# A multi-channel TMS system enabling accurate stimulus orientation control during concurrent ultra-high-field MRI for preclinical applications

**DOI:** 10.1101/2023.08.10.552401

**Authors:** Victor H. Souza, Heikki Sinisalo, Juuso T. Korhonen, Jaakko Paasonen, Mikko Nyrhinen, Jaakko O. Nieminen, Maria Koponen, Mikko Kettunen, Olli Gröhn, Risto J. Ilmoniemi

## Abstract

Monitoring cortical responses to neuromodulation protocols on preclinical models can elucidate fundamental mechanisms of brain function. Concurrent brain stimulation and imaging is challenging, usually compromising spatiotemporal resolution, accuracy, and versatility. Here we report a non-invasive brain stimulation system with electronic control of neuromodulation parameters in a 9.4-T magnetic resonance imaging (MRI) environment. In the imaging scanner, transcranial magnetic stimulation is delivered with a set of two coils and the MRI signals are recorded with a radiofrequency coil. The coil set provides millisecond-scale electronic control of the stimulus orientation with 1° resolution. Without physically rotating the coils, we evoked orientation-specific muscle responses after cortical stimulation on an anesthetized rat. We show that the stimulation pulses do not affect the anatomical imaging quality, and imaging signals are disrupted only if recorded before 10 ms after pulse delivery. Concurrent electronically targeted brain stimulation and neuroimaging sets the stage for the causal investigation of whole-brain network functions, endorsing more efficient treatment protocols.

## Introduction

Brain processes and diseases engage multiple cortical and subcortical regions that interact in a timely and orchestrated manner, forming complex interconnected networks (Lynn and Bassett, 2019). Instead of focusing on specific areas of the cortex, we need to understand brain function at the network level (Betzel and Bassett, 2017; Gordon et al., 2023; Siddiqi et al., 2022; Yang et al., 2021). Brain functions that span multiple cortical regions are observed across species, which makes preclinical experiments relevant for investigating biomarkers and the effects of treatments in pharmacologically, surgically, and genetically induced disease models (Dawson et al., 2018; Nestler and Hyman, 2010; Surendrakumar et al., 2023; Tang et al., 2017, 2020). In this context, transcranial magnetic stimulation (TMS) offers a non-invasive way of evoking targeted brain activation, and in combination with functional neuroimaging, it presents a powerful tool for assessing cognition and behavior (Allen et al., 2007; Bergmann et al., 2021; Ilmoniemi et al., 1997; Siddiqi et al., 2022, 2021; Siebner et al., 2009) and for optimizing therapeutic outcomes (Cash et al., 2021; Siddiqi et al., 2023). While TMS can modulate neuronal activity through processes like long-term potentiation and depression (Funke, 2018; Lefaucheur et al., 2014; Tang et al., 2017), neuroimaging methods, such as positron emission tomography (PET) (El Arfani et al., 2017; Krieg et al., 2013; Parthoens et al., 2012; Paus et al., 1997; Tastevin et al., 2020; Wyckhuys et al., 2013), electroencephalography (EEG) (Ilmoniemi et al., 1997; Li et al., 2007; Massimini et al., 2005; Rosanova et al., 2009; Rotenberg et al., 2008), and functional magnetic resonance imaging (fMRI) (Bergmann et al., 2021, 2016), provide spatial and temporal information on the underlying brain activity. EEG has high temporal resolution but offers limited spatial specificity, especially with rodents, and while PET provides higher spatial specificity than EEG, it requires radioactive tracers, hindering large-population studies. In turn, modern ultra-high-field MRI scanners offer superior spatial resolution compared to PET and EEG, enabling non-invasive network-level investigations of healthy and diseased brains (Bergmann et al., 2021; Gordon et al., 2023; Siebner et al., 2009; Tik et al., 2023).

Combining TMS with fMRI in a preclinical setting poses significant technical challenges (Mizutani-Tiebel et al., 2022; Seewoo et al., 2018). For instance, the restricted space prevents accurately targeting the cortical stimulation inside the scanner bore. It is also extremely difficult to change the targeting parameters during an experiment, and for instance, studies requiring subsequent pulses on different loci or in different orientations with millisecond-scale intervals are not possible with traditional instruments (Nieminen et al., 2019; Tugin et al., 2021). Moreover, the interaction between the high static magnetic field and stimulation current in the TMS coil windings leads to high mechanical stress, causing fractures (see Supplementary Video) to the coil structure and potentially exposing the subject to hazards (Bergmann et al., 2021; Crowther et al., 2013). Recently, the development of multi-locus TMS (mTMS) has enabled electronic control of the induced electric field orientation and location in the human brain with resolutions exceeding 1° and 1 mm, respectively (Nieminen et al., 2022b; Souza et al., 2022). This makes mTMS suitable for automated and highly accurate stimulus targeting inside the scanner bore. However, existing mTMS–MRI instrumentation has not been designed with mechanical properties and dimensions applicable to preclinical ultra-high-field MRI settings.

Here we aimed to develop an MRI-compatible mTMS system tailored for preclinical research. Also, our goal was to develop a system that allows for real-time electronic control of the orientation and intensity of applied brain stimulation within the MRI scanner, seamlessly integrated with imaging sequences. This system would offer unique features and improvements over existing devices for humans (Bestmann et al., 2003; Navarro de Lara et al., 2017), leading to significant breakthroughs in brain studies.

## Results

### MRI-compatible mTMS system design

For brain stimulation and imaging inside the shielded room, we designed, deployed, and validated an mTMS system capable of operating concurrently with a 9.4-T MRI scanner. The main components of the mTMS system are power electronics, cabinet structure, and a multi-coil transducer, illustrated in Fig. 1. This mTMS power electronics is a smaller, two-channel variant of our recently developed 6-channel mTMS system for human applications; details of the system architecture are described in our previous work (Nieminen et al., 2022b). Currently, the system consists of two independent channels (capable of controlling two coils), with support for adding a third channel.

**Fig. 1.**
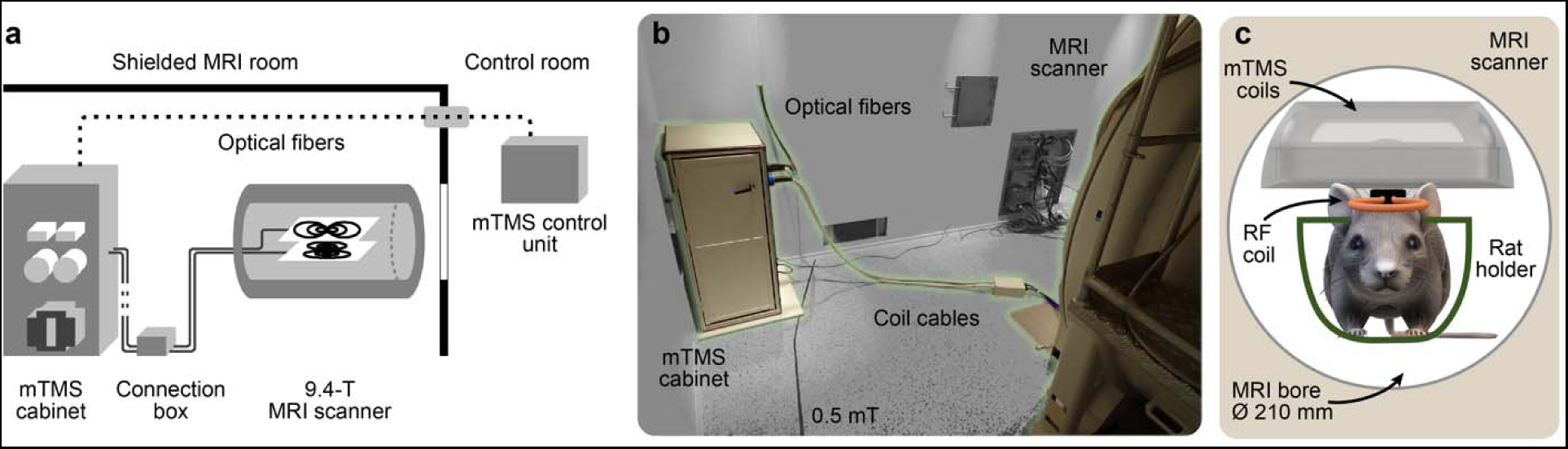
The mTMS–MRI system. **a**, Schematic of the mTMS system installation in the MRI shielded room. The optical fibers originating from the cabinet connect to the control unit inside the MRI control room. The mTMS transducer is placed in the MRI scanner bore and connects to the mTMS cabinet via two connection boxes, one inside the MRI bore (not visible in the photo or the schematic). **b**, Photo of the mTMS installation inside the MRI shielded room. **c**, Illustration of the stimulation and recording coils inside the MRI bore with a 210-mm diameter. The radiofrequency (RF) coil is used for recording the MRI signals and is attached to the bottom of the case with the mTMS coils. A custom holder supports the animal and the coils (see Fig 3a).

Each channel has an H-bridge module (Koponen et al., 2018, 2017) (maximum voltage 1500 V), which drives a current pulse in a single coil in an mTMS transducer. To minimize the electrical interference during mTMS–MRI recordings, the charging of the high-voltage capacitors is interleaved with the MRI acquisitions. All the components were designed to comply with MRI safety and compatibility (see Methods), with minimal ferromagnetic parts from structural components to printed circuit boards.

Compared to our six-channel mTMS device, a notable change is the split installation (Fig. 1a–b) with the power electronics cabinet fixed to a corner inside the shielded room outside the 0.5-mT region, and the control unit placed inside the operating room. This separation maintains a noise-free environment inside the shielded MRI room while enabling full control of the system and the MRI computer. With the power electronics inside the shielded room, we reduce the length of the cables connecting to the transducer, minimizing undesirable inductance and resistance critical for the system’s operation. The mTMS system supports connection to digital temperature sensors that can be embedded into the transducer for monitoring its temperature, and the signal path is protected against coil-induced overvoltage. The system also supports the electronic recognition of transducers via microchips with unique identifiers.

### mTMS transducer

To electronically manipulate the induced electric field (E-field) orientation in the rat’s cortex, we designed and built a 2-coil transducer. The winding paths for the coils were generated with a minimum-energy optimization method (Koponen et al., 2018) with wire density constraints (Rissanen et al., 2023) implemented in MATLAB 2022a. The coil winding paths were machined in polycarbonate plates due to their excellent mechanical properties (75 MPa tensile strength and 2.4 GPa Young’s modulus) necessary to withstand the high stress generated when applying TMS pulses with kiloampere-level currents inside an MRI bore with a 9.4-T static magnetic field. Each coil was wound with copper litz wire in the grooves of the coil former (Fig. 2a) and crimped to the connection box placed inside the scanner’s bore. The transducer plates are potted with a silicone-based elastomer so that the 5-mm-thick top of the top former serves as a cover. The coil assembly is housed inside a custom case machined from polycarbonate with a 1-mm-thick bottom for additional mechanical strength and safety (Fig. 2a).

**Fig. 2.**
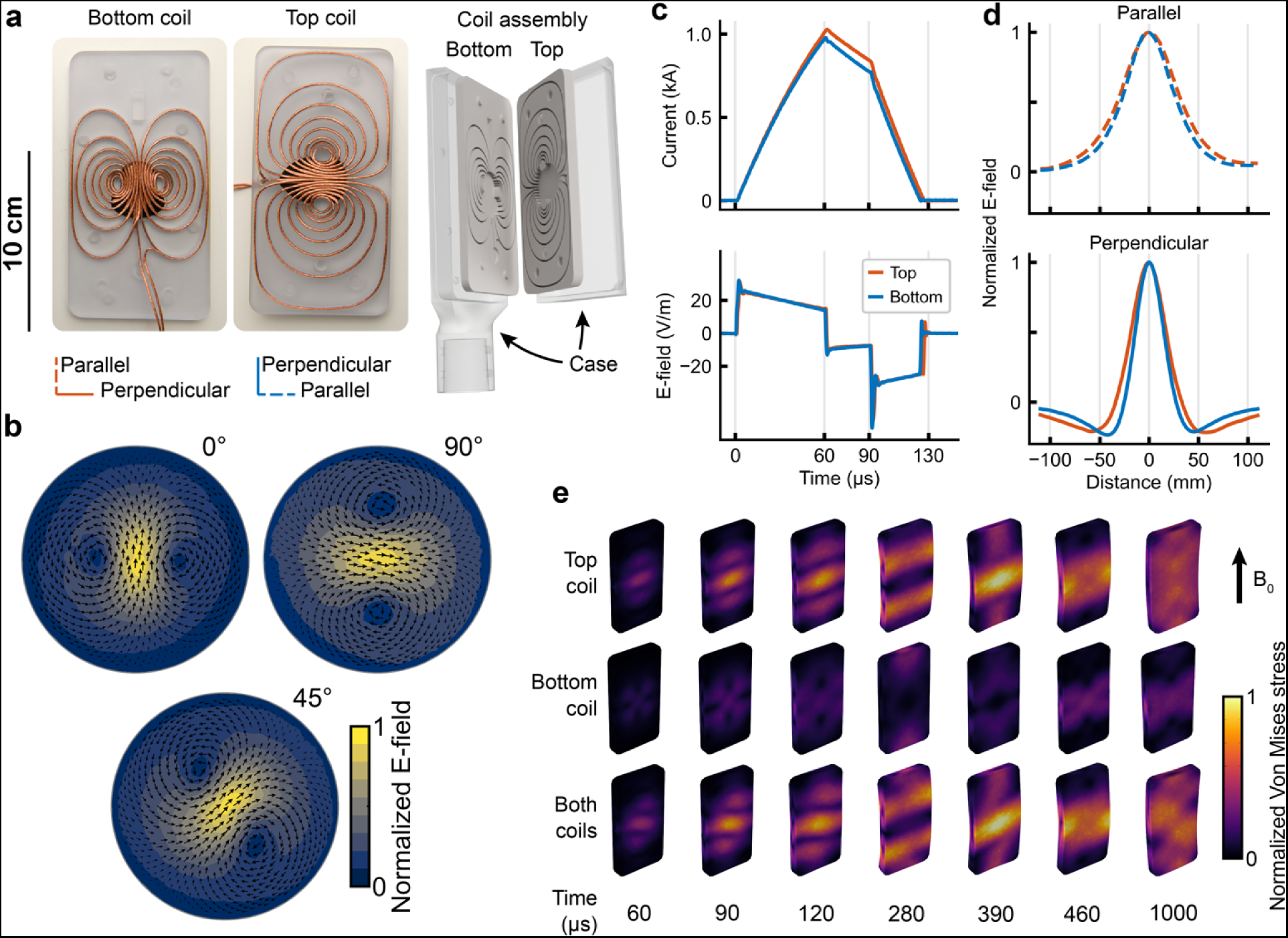
The mTMS transducer and measured E-fields. **a**, The two figure-of-eight coils wound with litz wire in polycarbonate formers and a 3D model of the transducer assembly with the protective case. **b**, E-field spatial distribution measured with the TMS characterizer on a 70-mm radius spherical head model. The 0°, 90°, and 45° correspond to the E-field orientation induced with the bottom coil, top coil, and both coils together, respectively. **c**, Trapezoidal monophasic current (upper panel) pulse and its induced E-field waveform (lower panel) for the bottom and top coils inducing an E-field on a 70-mm radius spherical head model. **d**, E-field profiles as indicated in (a) and measured with the same setup as in (b). Based on these profiles, we estimated the induced E-field focality in the direction parallel and perpendicular to the peak E-field. **e**, Simulated stress distributions on the coil plates inside the MRI bore with a 9.4-T static magnetic field (B_0_) when mTMS pulses were applied to each coil separately (bottom and top) or to both coils simultaneously. The time is relative to the mTMS pulse onset as in the current waveform illustrated in (c).

After assembling the 2-coil mTMS transducer, we calibrated the electronic control of the induced E-field orientation. The resulting E-field spatial distributions for different E-field orientations: 0° (bottom coil), 45° (both coils), and 90° (top coil) are shown in Fig. 2b. The E-fields were measured and calibrated over a spherical cortex model with 70-mm radius using our TMS characterizer (Nieminen et al., 2015), and with monophasic pulses delivered by the mTMS power electronics and had phases lasting 60 (rise) and 30 µs (hold), and a falling phase of 32.5 and 35.2 µs for the bottom and top coils, respectively (Fig. 2c). The bottom and top coils required capacitor voltages of 1030 and 1414 V, respectively, to induce an E-field of 70 V/m on the 70-mm radius cortex. The equivalent E-field on a rat cortex with 13.4-mm radius is 57 and 45 V/m for the bottom and top coil, respectively. The ratio between the E-fields on the 70- and 13.4-mm cortices was computed with the surface current distributions of the coil models obtained during the winding path optimization (see Methods). The induced E-field focality was computed as the full-width at ∼71% (1⁄√2) from the maximum of the perpendicular and parallel E-field profiles (Nieminen et al., 2015), shown in Fig. 2d. The bottom and top coil focalities in the direction perpendicular to the peak E-field were 21.4 and 25.7 mm, respectively; in the parallel direction, they were 37.8 and 42.9 mm, respectively.

To assess the stress distribution and mechanical durability of the coil formers, we performed finite element simulations in COMSOL Multiphysics® 6.0 (COMSOL AB, Sweden) using a physically accurate 3D model of the transducer copper wiring, polycarbonate plates, and on a uniform magnetic field of 9.4 T representing the MRI bore. The stress was considerably higher for the top coil or with both coils fired together than for the bottom coil alone. The simulated peak stress values with a monophasic stimulation pulse (rise: 60 µs, hold: 30 µs, fall: 32.5 for the bottom and 35.2 µs for the top coil; see Fig. 2c) at 100% of maximum stimulator output (MSO) were 31.5 MPa, 70.9 MPa, and 71.4 MPa for the bottom coil, top coil, and both coils together, respectively. The change in stress distribution over time suggests that the coil plates vibrate and deform and may eventually break. We took the results into account in the designs of the coil formers by increasing the thickness of the top plate compared to the bottom and by reinforcing the central region with a fluoroelastomer (black surface below the coil wires in Fig. 2a) to dampen the impact from the wires.

### Interplay between mTMS and MRI

To ensure the compatibility of the mTMS device and the MRI sequences, we characterized the mTMS transducer effect on the radiofrequency (RF) coil’s resonance frequency and the presence of imaging artifacts due to the stimulation pulses. The MRI signal was recorded with a custom-made 400-MHz single-loop transceiver surface coil (Neos Biotec SL, Spain). The proximity of the mTMS transducer shifted the RF coil’s resonance frequency by 7 MHz, which was considered while optimizing the tuning and matching range of the RF coil. The mTMS system did not induce visually observable artifacts in the structural MRI. The mTMS transducer was not visible in the spin echo images, but it was faintly visible in the images obtained with the gradient echo sequence (Fig. 3b). The free induction decay (FID) signal was distorted up to 3.0 and 6.0 ms after applying mTMS pulses with 8% and 25% MSO, respectively. The 100% MSO corresponds to a 1500-V pulse driven by each channel on the mTMS power electronics. The top coil alone (−90° orientation) caused the longest period of signal corruption in the FID power spectrum (up to 6.0 ms at 25% MSO pulse) while utilizing two coils simultaneously (e.g., a pulse in 45° orientation) caused the shortest period of signal corruption (up to 4.0 ms) (Fig. 3C–D). Additionally, we tested whether the powering of the mTMS system or connecting the mTMS transducer cables induced noise to the MRI’s FID signal with 10 kHz and 208 kHz bandwidths without applying any mTMS pulses. The presence of active but non-stimulating mTMS electronics affected the FID signal only minimally with 10 kHz (difference in AUC < 0.001 %) or 208 kHz (difference in AUC < 0.01 %) bandwidths.

**Fig. 3.**
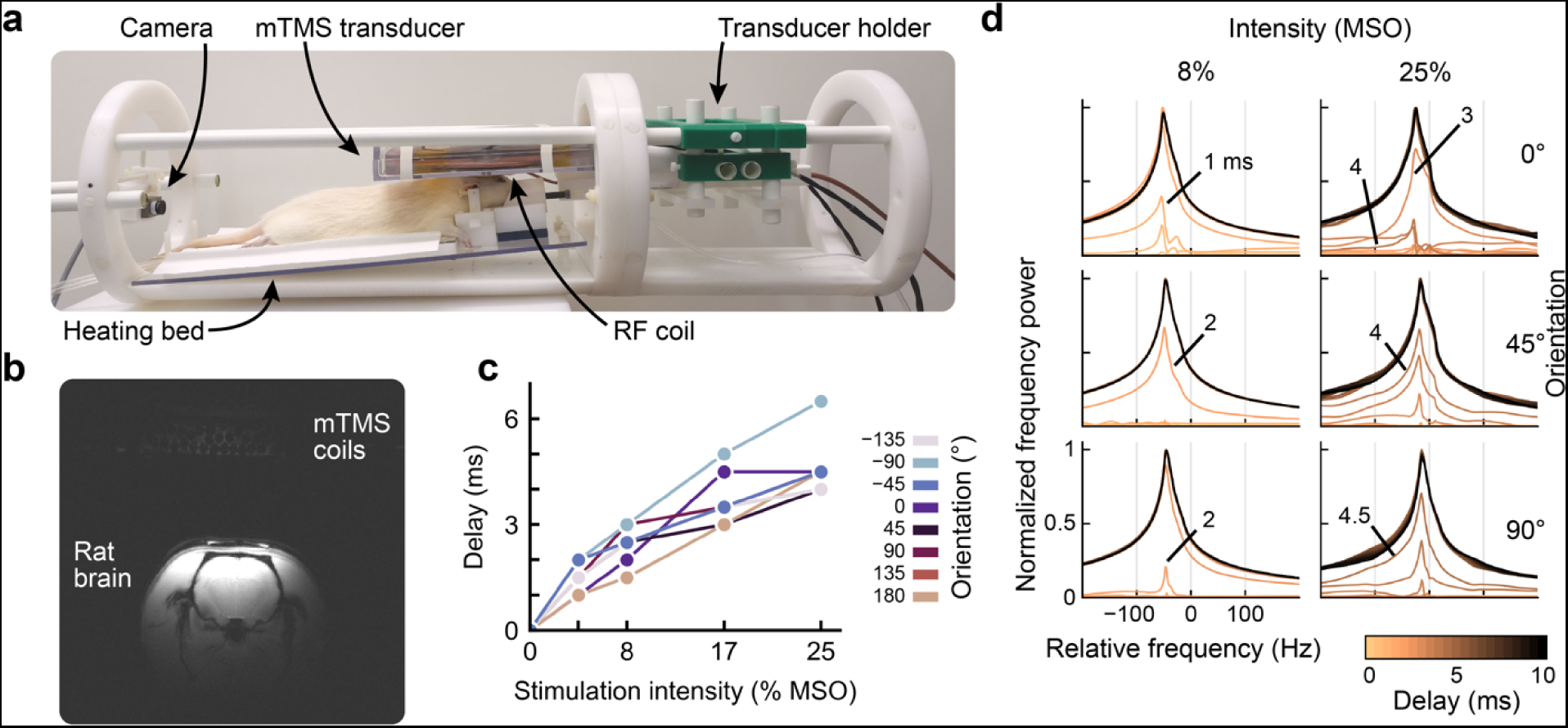
Effect of mTMS pulse on the MRI signal. **a**, Experimental setup for positioning the mTMS transducer and the fMRI-RF coil over the rat’s head inside the MRI bore. **b**, MRI taken with the mTMS transducer on top of the rat brain. The image brightness was increased to make the mTMS coil windings visible. **c**, Minimum delay between the mTMS pulse and the RF excitation pulse to obtain an artifact-free FID signal at different E-field orientations. Stronger intensities require longer delays which also depend on the induced E-field orientation. Stimulation intensities are measured as % of the maximum stimulator output (MSO). **d**, MRI signal normalized power spectra recorded with multiple mTMS pulse intensities, orientations, and delays relative to the RF pulse. The black line represents the spectrum without an mTMS pulse. The numbers indicate the longest millisecond delay with a corrupted power spectrum. At 0° orientation and with a stimulation intensity of 25 % MSO, the 4-ms delay showed a more corrupted spectrum than a 3-ms delay.

### Acoustic noise generated by the mTMS pulses

We measured the sound pressure levels (SPL) with the open end of a 6.5-m long non-elastic tube in five locations: 4 cm below the mTMS transducer center and at a 1-m distance (far field; see Methods), both measurements with the transducer inside and outside the MRI. The fifth location was with the mTMS transducer inside the MRI bore and the tube’s open end in the MRI operation room (Fig. 2). The SPL levels were computed without weighting, i.e., Z-weighting (Acoustical Society of America, 2014). The peak SPL measured outside the MRI bore (without the B_0_ field) was 142 dB(Z) at a 180° stimulus orientation and with 100% MSO stimulation intensity (Fig. 4). The effect of orientation across all stimulation intensities was 9 dB(Z) on average, 0° being the loudest and −90° the quietest. Inside the bore, the highest peak SPL was 167 dB(Z), measured with a 0° pulse orientation (earplug as a dampener) using 80% MSO stimulus intensity. On average, the static magnetic field increased the SPL by 44 dB(Z), with stimulation intensities of 20% and 40% MSO. We also measured the SPL inside the control room with the mTMS transducer inside the bore using a 20%-MSO stimulation intensity. In the control room, SPLs were below 85 dB(Z), and it was impossible to distinguish them from the background noise with our measurement system. Thus, there is at least 60 dB of noise reduction from the scanner center to the control room. With the same intensity (20% MSO), we measured the effect of sound-insulating polyurethane foam (HiLo-N40, t.akustik, Germany) inside the MRI scanner and with the measurement tube at a 4-cm distance from the mTMS transducer bottom. The foam covered the MRI bore walls to dampen the sound wave reflections. On average, the sound-insulating foam lowered the peak SPL measured below the coil by 4 dB(Z). At a distance of 1 m, the peak SPLs inside and outside the bore were attenuated by 20 dB(Z) and 23 dB(Z) on average, respectively. Fig. 4 shows the SPLs for mTMS pulses inside and outside the MRI for all tested stimulus intensities and orientations.

**Fig. 4.**
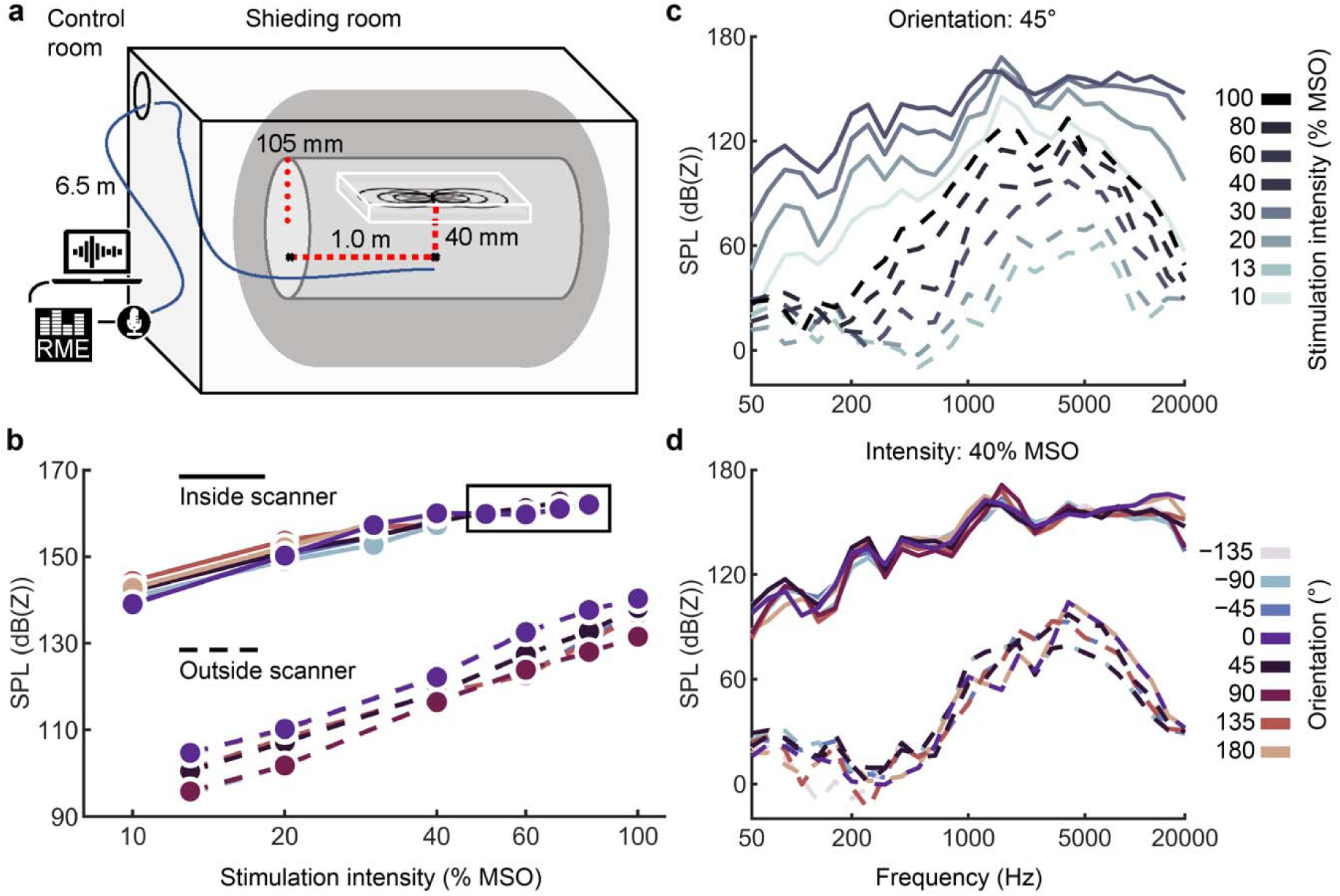
Acoustic noise from the mTMS pulses. **a**, The acoustic noise measurement setup inside the MRI bore with the open end of the measurement tube pointing towards the transducer and placed inside the MRI bore in the two positions indicated by the black dots connected with red dashed lines: 1) 40 mm below the bottom of the mTMS transducer’s center for near-field measurements and 2) at a 1-m distance from the first measurement point for far-field measurements. The tube orientation and position in the image are for illustration purposes only. **b**, peak SPLs measured outside and inside the MRI scanner with the open end of the tube at 4 cm below the transducer with different stimulation intensities and orientations. The measurements inside the square were performed using an earplug as a dampener at the open end of the tube. **c**, 1/3 octave spectra of the acoustic noise from mTMS pulses at 45° orientation (bottom and top coils together) for different intensities. **d**, 1/3 octave spectra for mTMS pulses at 40% MSO with varying stimulus orientations. In all charts, dashed and solid lines indicate measurements outside and inside the MRI scanner, respectively.

### Effects of stimulus orientation on motor response

To verify that the mTMS transducer stimulated the rat’s brain, we measured the effect of mTMS stimulus orientation on the amplitude and latency of motor-evoked potentials (MEPs) in an anesthetized rat outside the MRI bore. We applied five single with 110% of the resting motor threshold (RMT) in each of eight stimulus orientations (from −135° to 180° in steps of 45°) over the motor hotspot at the left and right brain hemispheres while recording electromyography (EMG) from ipsi- and contralateral biceps brachii muscles. The RMT was 73% MSO at 0° orientation. Fig. 5 shows that the MEP amplitude considerably depended on the stimulated brain hemisphere and the E-field orientation. On the right brain hemisphere, contralateral MEPs with the highest amplitudes were evoked at −90°, −45°, 90°, and 135°, while no MEPs were detected at the remaining orientations. In turn, the left hemisphere showed the highest contralateral MEP amplitudes for mTMS pulses at 45°, 90°, and −135°, with no MEPs at −90°, −45°, and 180°. We detected ipsi- and contralateral MEPs in both hemispheres with latencies of about 10 ms.

**Fig. 5.**
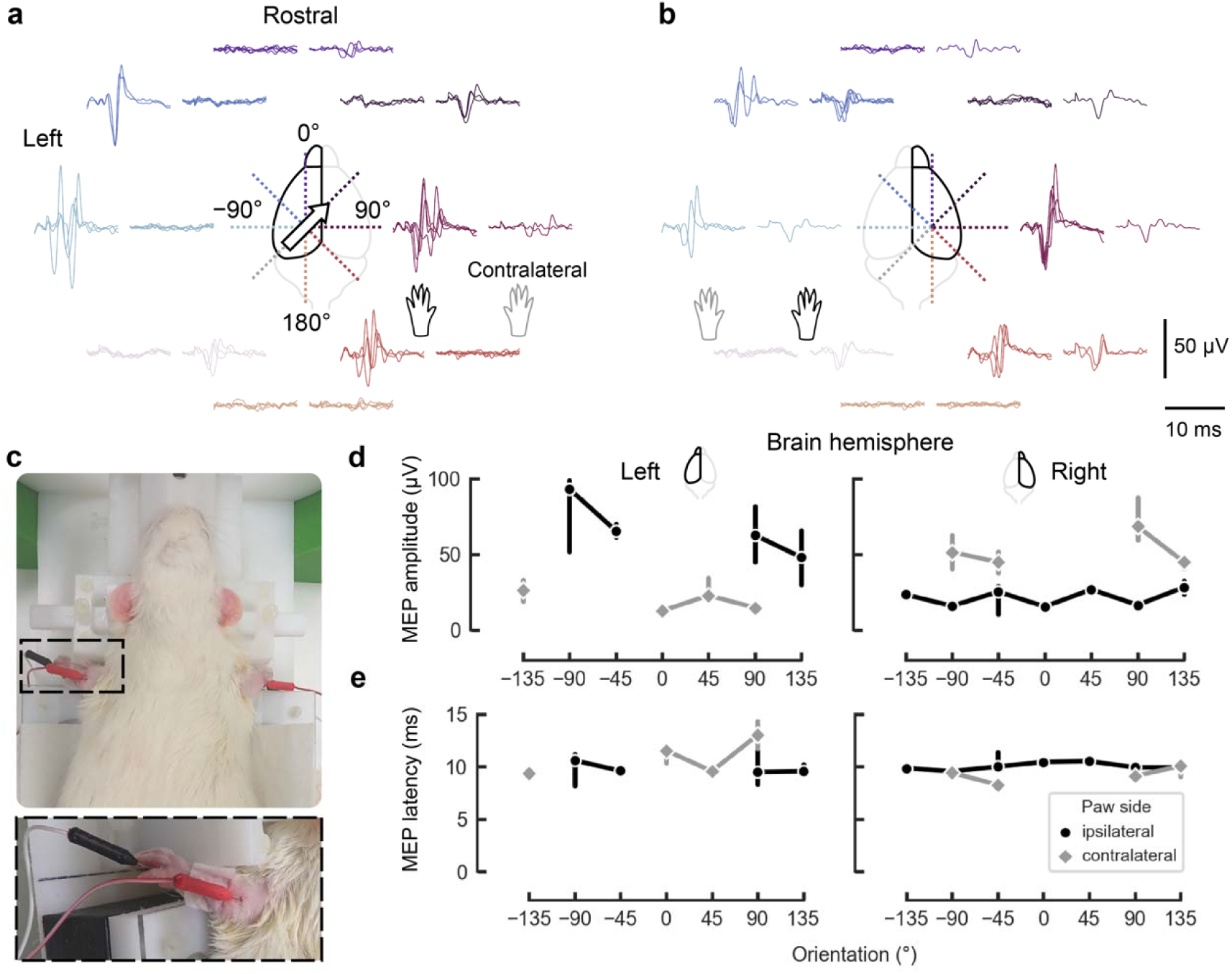
Effect of mTMS pulse orientation on MEPs. MEPs recorded from the rat’s left and right biceps brachii with mTMS pulses on the **a**, left and **b**, right rat brain hemispheres. MEPs were recorded with a stimulus intensity of 80% MSO (110% of resting motor threshold). **c**, Photos depicting the rat placement on the holder and the needle electrode placement on the right and left front paws. The dashed square highlights the electrode placement. **d**, MEP amplitude from ipsilateral (black circle) and contralateral (gray diamond) paws after stimulation of the rat’s left and right brain hemispheres with stimulus orientations varying from −135° to 180°. Markers represent the medians across trials and the error bars represent the 95% confidence interval. **e**, Latency for the corresponding MEP amplitudes. Missing latency values result from MEPs with zero amplitude or trials rejected during signal preprocessing. The biggest contralateral MEPs were recorded with mTMS pulses on the right brain hemisphere (b) and with the E-field orientations 45°, 90°, 135°, and −90°.

## Discussion

We introduce and demonstrate the first in-situ mTMS–MRI setup for concurrent stimulation and imaging of small animals in an ultra-high-field MRI scanner (static field 9.4 T). The electronic control of TMS stimulus parameters with the transducer inside the MRI bore solves important limitations of concurrent TMS–MRI applications. For instance, with specialized multi-coil transducers (Rissanen et al., 2023), distinct cortical areas can be subsequently targeted within milliseconds with interleaved imaging sequences, allowing causal assessment of network brain functions (Siddiqi et al., 2022). With electronic targeting, we also minimize the need to reposition the stimulating coil physically. Such repositioning requires the operator to remove the experimental setup from the bore, which is time-consuming and prone to errors.

The mTMS–MRI installation with the cabinet inside the shielded room and the control unit outside provides significant advantages compared to existing systems (Bestmann et al., 2003; Navarro de Lara et al., 2017). This installation minimizes the length of the transducer cabling, whereas commercial MRI-compatible TMS systems typically utilize extra-long coil cables and introduce the coil into the MRI room via a waveguide combined with a low-pass filter. This has been reported to result in a loss of up to 15% in stimulation strength (Bergmann et al., 2021). The stimulation intensity is a critical limiting factor in TMS in preclinical applications due to the reduced electromagnetic coupling between the animal’s brain and the stimulation coils. Thus, even minor improvements in stimulation intensity might enable a broader range of stimulation protocols. Another crucial improvement is that the computer that controls the mTMS device from the MRI control room allows flexible programming of mTMS pulse waveforms and sequence protocols. This feature eliminates the need for the operator to frequently enter and exit the shielded room during experimental sessions, reducing exposure to potential health risks, such as electric hazards and high acoustic noise levels (discussed below).

A unique characteristic of our system is the electronic control of the E-field orientation in the rat cortex enabled by the 2-coil mTMS transducer inside the MRI bore. We combined rigid polycarbonate plates with a layer of fluoroelastomer and potting with a special silicone for improved impact absorption. This layered design withstood stimulation intensities up to 80% of MSO in all orientations. This is a significant improvement compared to our first prototypes manufactured with only polyoxymethylene (POM) and potted with epoxy, which fractured at about 40% MSO. Further research is still required to achieve more durable, efficient, quiet, and, if desired, more focal stimulation coils. In applications not requiring high focality, it might be best to trade the focality fully for efficiency (Nurmi et al., 2021), as such coils can evoke reliable MEPs in rats with relaxed durability requirements (Nieminen et al., 2022a). Importantly, the multi-channel TMS system is flexible and can be utilized with multiple coil array combinations designed for specific purposes, such as precise electronic cortical mapping (Nieminen et al., 2022b; Rissanen et al., 2023).

Our phantom measurements indicate that the MRI-compatible mTMS system is suitable for conventional preclinical MRI studies. With charged mTMS pulse capacitors, the stimulation coils inside the bore induced minimal changes in the FID signal, indicating that the coil cables did introduce high-frequency noise, and the potential small leakage current from the high-voltage capacitors had a minimal effect. We did not observe any significant image distortions, and there was only a small signal originating from the mTMS transducer in commonly used gradient and spin echo sequences. In addition, only a 6-ms delay was required after the mTMS pulse to have a negligible effect on the FID data. We should note that our mTMS–MRI delay measurements were conducted only up to 25% MSO. However, based on a logarithm extrapolation, the required delays are within 10–20 ms for higher voltages (e.g., 67% MSO). This delay range is suitable for interleaved mTMS–MRI recordings and lower than the minimum reported in previous studies, in which 50–100 ms have been suggested to avoid TMS artifacts in echo planar images (Bestmann et al., 2003; Navarro de Lara et al., 2017). An evident limitation of our study is that no TMS-evoked functional MRI signals were measured from the rat’s brain. Such measurement falls outside this study’s scope and could potentially lead to unnecessary complexities in interpreting the neurophysiological effects of brain stimulation that would be better explored in a future study.

The SPL increased logarithmically with increased stimulation intensity, and there were small differences depending on the stimulation orientations. At a 1-m distance from the coil, the acoustic noise levels were about 20–23 dB lower than at a 4-cm distance, so proper safety measures must be in place to prevent accidental mTMS pulses when personnel are working close to the bore, for example, during the preparation of the animal for the experiment. The safety limit for the peak SPL for humans (140 dB(C), computed with C-weighting) was exceeded during stimulation both outside and inside the MRI bore. Still, it remained at safe levels in the operation room. In general, laboratory animals exposed to continuous sound suffer from the same reactions to noise, ranging from auditory effects like temporary or permanent threshold shifts and tinnitus to non-auditory effects emerging from noise-induced stress, such as elevated heart rate and sleeping problems (Lauer et al., 2012; Turner et al., 2007). Experiments with mTMS in an ultra-high-field MRI should be performed with anesthetized animals, as in this study, carefully considering the animal safety regarding exposure to impulse sounds.

As a proof of concept of the electronic control of stimulus orientation, we demonstrated that the MEPs considerably depended on the E-field orientation. The corresponding MEP latencies (approximately 10 ms) are akin to those reported in previous studies (Boonzaier et al., 2020; Nieminen et al., 2022a; Rotenberg et al., 2010). Interestingly, only specific transducer locations and orientations generated MEPs despite the relatively broad induced E-fields (20–30 mm range), which, in principle, would stimulate large cortical areas. This is further supported by the similarity of the MEPs across different orientations evoked by stimulation of both right and left hemispheres (see Fig. 7A–B), as it is likely that both transducer locations stimulated the same cortical region due to the widespread E-field patterns on the rat’s cortical surface (Koponen et al., 2020). Such high TMS spatial and orientation specificity could be due to a localized motor cortical representation by neuronal ensembles aligned in laminar and columnar structures (Kätzel et al., 2011), as extensively studied in humans (Fox et al., 2004; Souza et al., 2022; Tardelli et al., 2022; Weise et al., 2023a), and further supported by the stimulation of subcortical neuronal pathways with deep brain stimulation electrodes (Lehto et al., 2020; Slopsema et al., 2018). Notably, simulations on morphologically realistic neurons corroborate the directional sensitivity of pyramidal neurons to electromagnetic stimuli (Aberra et al., 2020; Weise et al., 2023b; Wu et al., 2016). In addition, simulations also indicated an absolute E-field intensity to depolarize pyramidal neurons similar to our estimated values on a spherical rat cortex (Wu et al., 2016).

In conclusion, our system enables concurrent non-invasive neuroimaging and brain stimulation applications in a preclinical setting. Our mTMS coils were designed to provide fast electronic control of stimulus orientation as a new feature for rat brain stimulation with concurrent MRI. Due to the flexibility of our mTMS power electronics and coil design algorithms, we open a possibility for future designs with critical improvements in stimulus focality, which can elucidate better the underlying physiological and anatomical processes on the orientation sensitivity of the rat brain motor representations. This setup can be utilized to create and test a wide range of protocols to study whole-brain and network effects of neuromodulation (Bergmann et al., 2021; Mengotti et al., 2022; Tik et al., 2023; Yang et al., 2021) and treatments for neurological disorders (Dawson et al., 2018; Seewoo et al., 2018; Siddiqi et al., 2022).

## Methods

### mTMS cabinet

The device cabinet was custom-made from aluminum profiles and panels, non-magnetic stainless-steel fasteners (A2 and A4 grades), and polyoxymethylene plastic mounting plates. The only major magnetizable component in the system that could not be placed outside the shielded room was a 2000 VA isolation transformer. Whenever possible, internal components were fixed to the cabinet’s structure. The metal structure of the cabinet is earthed with a separate cable, which remains connected even when the power cable is disconnected. The earthing follows a “star-ground” design, thus eliminating potential ground loops. The electronics are isolated by an isolation transformer (resistance to ground >2 MΩ), providing added user safety. There are no electrical connections between the power electronics and operator control cabinets. For thermal management, three 120 mm diameter fans push out air from the top of the cabinet, and the replacement air is mainly taken in from around the cabinet’s front door, providing air flow from front to back and bottom to top. A custom-made door handle, lock, and non-magnetic copper keys were manufactured to prevent unauthorized access to the cabinet. The power electronics were designed to default into a safe state during the sudden absence of power. The cabinet also includes an electronic safety system, which immediately discharges the pulse capacitors if the cabinet door is opened. The system also contains an emergency stop button.

In the split installation (Fig. 1), the control unit communicates with the power electronics via fiberoptic links fed through a waveguide. The power electronics are controlled by a reprogrammable I/O platform (PXIe-7846R, National Instruments, Corp., USA) housing a field-programmable gate array (FPGA, Kintex-7 160T, Xilinx, Inc., USA) running at a clock frequency of 40 MHz. The mTMS device can be operated through a graphical user interface programmed with LabView 2021 (National Instruments) or via an application programming interface enabling customized solutions, for instance, in Python or MATLAB (The MathWorks, Inc., USA).

### mTMS transducer design

First, we modeled a small commercial figure-of-eight coil (MC-B35, MagVenture A/S, Denmark) on a triangular mesh (2568 vertices) located 15 mm away from the cortical surface (13.7-mm-radius sphere; 2562-vertex triangular mesh). With the commercial coil model, we computed the induced E-field distribution for two coil orientations (0° and 90°). For both orientations, we computed in a rectangular plane section (19-cm long and 9.5-cm wide; 1953-vertex triangular mesh) the corresponding minimum-energy surface current density that induces an E-field distribution with similar focality and intensity than the commercial coil model. We defined the dimensions of the rectangular plane section to fit inside a small gradient coil set (ø 11–12 cm) typically used in preclinical small-bore MRI systems. For the 0° and 90° coil orientations, we performed the optimization with the rectangular plane located at 15 and 20 mm from the cortical surface, respectively. Then, we decomposed the optimized surface current densities with singular value decomposition and discretized the coil winding paths from each surface current density in 14 isolines. This resulted in two orthogonal figure-of-eight coils (seven turns in each wing). The coils were wound with 1.7-mm diameter litz wire (3-layer Mylar coating; Rudolf Pack GmbH & Co. KG, Germany) and potted with a 2-component silicone-based elastomer (SYLGARD® 184, Dow, Inc., USA).

To enable fast prototyping of different transducer designs, we designed a connection box inside the MRI bore that allows simple, fast, and safe assembly and connection of the coil cables to the cabinet with threaded copper rods and plastic plates for electrical insulation. In the connection box, the coil cables are connected with lugs to a 2.5-m-long low-inductance TMS coil cable (The Magstim Company, Ltd., UK) that is connected in series with another 2.5-m-long low-inductance TMS cable (Nexstim, Plc., Finland) connecting to the mTMS power cabinet through a medical-grade connector (Nexstim).

### mTMS transducer manufacturing and calibration

To manufacture the transducer, we designed two plates (bottom and top coils) and a case in SolidWorks 2018 (Dassault Systèmes SA, France). The bottom former thickness is 7.0 mm (including a 2.0-mm-thick bottom), and the top former is 10.0 mm thick (including a 5.0-mm-thick top). The bottom thickness corresponds to the material thickness below/above the wire grooves. The inner base of the outer case contains small studs, and the bottom coil plate has matching cavities. While the transducer is resting on the studs, a 1-mm air gap is present between the enclosure and the coil plates, reducing the mechanical coupling between them and providing an additional safety barrier in case of fractures in the coil plates.

When calibrating the transducer, we measured the E-field distributions at 1000 points over a spherical human cortex model with a 70-mm radius using a TMS characterizer (Nieminen et al., 2015). The center of the transducer case bottom was 85 mm from the center of the spherical head model. We computed the E-field intensity as its average norm on a trapezoidal monophasic current waveform’s rising phase (60-µs long). For estimating the absolute E-fields measured with the TMS characterizer on the spherical rat cortex with a 13.7-mm radius, we computed the E-fields induced on a 70-mm radius spherical cortex model by the decomposed surface current distribution described in the mTMS transducer design. The coils’ self-inductances were measured with an LCR meter (1-kHz reference frequency; ELC-130, Escort Instruments Corp., Taiwan) and the resistances with a 4-wire measurement setup using a benchtop multimeter (HP 34401A; Hewlett-Packard Company, USA). The top coil (16.7 µH, 133.0 mΩ) required approximately 37% more voltage than the bottom coil (12.7 µH, 121.82 mΩ) to induce the same E-field measured with the TMS characterizer. The duration of the trapezoidal monophasic pulse waveform was customized for each coil based on measurements with a Rogowski probe (CWT 60B, Power Electronic Measurements, Ltd., UK) connected to an oscilloscope (InfiniiVision MSOX3034T, Keysight Technologies, Inc., USA) to ensure that no current was left circulating in the system after a pulse.

### Finite element modeling

The coil plates were placed inside a cylindrical air domain with a uniform magnetic field of 9.4 T, representing the MRI bore. The coil plates were fixed in space by applying a spring boundary condition to one of the plates’ ends with a spring constant per unit volume of 1 N/m^3^. The pure copper and polycarbonate material properties were obtained from the COMSOL material library and the Granta EduPack 2022 R1 database (ANSYS, Inc., USA), respectively.

### mTMS–MRI characterization

MRI measurements were done with a 9.4-T magnet (bore diameter 31 cm) interfaced with a DirectDRIVE console (Agilent Technologies, Inc., USA) and with a 21-cm-inner-diameter gradient coil set. During manufacturing, the tuning and matching range of the RF coil was optimized while it was centered at the bottom of the mTMS transducer case. We characterized the effect of mTMS pulses on the MRI signal excitation and acquisition in an experiment where we first applied a stimulation pulse and then acquired the free induction decay signal from a rat brain phantom after a single hard RF pulse. The RF pulse was delivered with a custom coil consisting of a 22-mm inner diameter open loop connected to a separate tuning and matching box with a 6-cm-long coaxial cable. The RF excitation pulse was given with a flip angle of 2° and a variable delay of 0–10 ms after the mTMS pulse (0.5 ms steps, 21 acquisitions with 2.5 s intervals) with a repetition time of 2500 ms. Slice selection was not used. For each mTMS–RF pulse delay, we tested eight stimulus orientations (−135°–180° in steps of 45°) with five intensities (0, 5, 8, 17, and 25% of maximum stimulator output).

### mTMS pulse acoustic noise

The sound pressure levels (SPL) were measured in five locations, as described in the Results section: 4 cm below the mTMS transducer center and at a 1-m distance (far field) inside and outside the MRI. The fifth location refers to the mTMS transducer inside the MRI bore and the tube’s open end in the MRI operation room. The beginning of the far field is given roughly by L^2^f⁄4v, where L is the largest dimension of the source, f is the frequency, and v is the speed of sound (Kinsler et al., 1999). For a TMS pulse, the far field starts at about 60 cm away from the coil center. In each location, we tested eight different E-field orientations (−135° to 180° in steps of 45°) and six stimulation intensities (13% and 20%–100% in steps of 20% of maximum stimulator output; MSO) outside and eight intensities (10%–80% in steps of 10% MSO) inside the MRI bore. For high intensities (> 40% MSO), we covered the open end of the tube with a foam earplug to prevent the SPL from saturating the audio interface input. To avoid potential damage to the mTMS transducer due to the high magnetic forces involved, we limited the maximum intensity inside the MRI bore to 80% MSO. The acoustic noise reduction was calculated by measuring the SPL for all E-field orientations at 20% MSO stimulation intensity with and without the earplug. The average reduction between these two conditions was added to the SPL measurements utilizing the earplug. The acoustic noise was measured with a high-performance microphone (MKE 2 P-C, Sennheiser electronic GmbH & Co. KG, Germany) firmly attached to the end of a non-elastic tube (Tress Nobel, 40 bar, length 6.46 m, diameter 6 mm; Tricoflex) and digitized with a high-quality audio interface (RME Babyface Pro, Audio AG, Germany) controlled with custom software written in MATLAB 2022a (Nyrhinen et al., 2023). To accurately represent the acoustic noise SPL and frequency spectrum, we calibrated our measurement setup to the frequency range of 20–20,000 Hz at the Aalto Acoustics Laboratory (Aalto University, Finland).

### Animal electrophysiological and imaging recordings

All animal procedures were approved by the Animal Experiment Board in Finland and conducted following the European Commission Directive 2010/63/EU guidelines. Neurophysiological experiments were performed with one adult male Wistar rat (400–600 g, RccHan®: WIST; Envigo RMS BV, Netherlands). Before the experiments, the rat was anesthetized with isoflurane (5% for induction, 2% for maintenance; Attane Vet 1000 mg/g, Piramal Critical Care BV, The Netherlands) in 30/70 O2/N2 carrier gas. Subsequently, anesthesia was switched to urethane (1250–1500 mg/kg i.p.; Sigma-Aldrich, Corp., USA). The rat was placed on a warm water circulation pad (37 °C; Corio CD, Germany) to maintain a normal body temperature. Sufficient depth of anesthesia was periodically confirmed by pinching the hind paw to observe the loss of limb contraction reflex.

To study whether the mTMS setup induces any observable image distortions or signals during MRI, we performed structural imaging with the mTMS transducer on top of an anesthetized rat in a custom-made holder. For that, MRI data were acquired with two pulse sequences: multi-slice gradient echo (repetition time 18 ms, echo time 3 ms, flip angle 20°, a field of view 60 mm × 60 mm, matrix size 128 × 128) and spin echo (repetition time 2000 ms, echo time 30 ms, flip angle 90°, field of view 50 mm × 50 mm, matrix size 256 × 256).

The head of the rat was fixed to our custom-made holder with ear and bite bars. The mTMS experiment started 30 minutes after the initial isoflurane anesthesia induction. EMG was recorded with disposable monopolar needle electrodes (SDN electrodes, stainless steel, Inomed GmbH, Germany) inserted into the belly of the biceps brachii muscle in the depilated forelimbs’ left and right paws (Boonzaier et al., 2020; Rotenberg et al., 2010). The muscle belly was determined by palpation of the extended forelimb. The references were placed between each paw’s second and third digits, and the ground was inserted in the tail’s base. EMG signals were digitized with a NeurOne Tesla (Bittium Biosignals Ltd., Finland) with 10 kHz sampling frequency and 2.5 kHz low-pass filtering.

To measure the effect of the E-field orientation on MEPs, we first mapped the scalp location showing the highest amplitude MEPs with the mTMS transducer center about 0.5 cm lateral to the bregma and with stimulation intensity set to 67% MSO. Stimulation intensity and transducer location were adjusted until we detected MEPs on the contralateral biceps brachii muscle with visible limb twitches. With the transducer location fixed on the hotspot, we searched for the resting motor threshold (RMT) as the minimum stimulation intensity evoking at least three out of six MEPs with amplitude over 10 µV. The interval between consecutive pulses was 4.0–5.0 s.

### MRI data analysis

To analyze the possible effects of charger and cable noise, and the effect of mTMS pulses on FID signal at different delays, a Fourier transform was performed, and the area under curve (AUC) was calculated from the obtained spectra. Each set of FID measurements was repeated 10–20 times. Potential image distortions and signals induced by the mTMS transducer were visually evaluated from the structural images processed with the Aedes software (aedes.uef.fi).

### Acoustic data analysis

The SPLs were calculated as the average of the maximum SPLs across three separate mTMS pulses. The frequency spectrum was measured with a time window spanning 3 ms before and 40 ms after the pulse. The window length was approximately the time for the sound waves to travel twice the length of the measurement tube. Data samples of equal length were selected to calculate the frequency responses of the background noise. The samples were cosine filtered using the Tukeywin function to reduce artifacts from cutting the signal. Then, the filtered signal was used to create the 1/3 octave spectra. Measurements with the earplug were left out from the 1/3 octave spectra because the earplug altered the frequency responses.

### MEP data analysis

The TMS pulse artifact was removed from the continuous EMG signal using the TMS-EEG Artefact Removal tool (Version 1.0; Mega Elektroniikka Oy, Finland). The exponential decay model coefficients were set to 0 and band-pass filtered (3^rd^-order Butterworth filter, 15 and 1000 Hz cut-off frequencies). The EMG signal was divided into epochs from −5 ms to 20 ms before and after the TMS pulse. MEPs were visually inspected, and epochs showing large artifacts, visible heartbeats, or having MEPs with peak-to-peak amplitude smaller than 10 µV were discarded from further analysis. In total, 33 trials were discarded from 80 recorded. We extracted the peak-to-peak amplitude from the remaining MEPs and manually annotated the onset latency.

The MRI, acoustic, and MEP data were processed and analyzed with custom MATLAB scripts.

## Acknowledgments

This project has received funding from the Jane and Aatos Erkko Foundation, the Academy of Finland (decisions No. 294625 and 349985), and the European Research Council (ERC Synergy) under the European Union’s Horizon 2020 research and innovation programme (ConnectToBrain; grant agreement No 810377). We acknowledge the computational resources provided by the Aalto Science-IT project.

## Data and code availability

The main data supporting the results of this study are available at OSF (https://osf.io/2qmzj/) and within the paper. Additional data and code can be requested from the corresponding author.

## References

Aberra AS, Wang B, Grill WM, Peterchev A V. 2020. Simulation of transcranial magnetic stimulation in head model with morphologically-realistic cortical neurons. Brain Stimul 13:175–189. doi:10.1016/j.brs.2019.10.002

Acoustical Society of America. 2014. American National Standard Electroacoustics - Sound level meters - Part 1: Specifications (a nationally adopted international standard) - ANSI/ASA S1.4- 2014/Part 1, IEC 61672-1:2013. ed, S1.4-2014/Part 1 / IEC 61672:1-2013. Melville, NY: Acoustical Society of America.

Allen EA, Pasley BN, Duong T, Freeman RD. 2007. Transcranial magnetic stimulation elicits coupled neural and hemodynamic consequences. Science (1979) 317:1918–1921. doi:10.1126/science.1146426

Bergmann TO, Karabanov A, Hartwigsen G, Thielscher A, Siebner HR. 2016. Combining non-invasive transcranial brain stimulation with neuroimaging and electrophysiology: Current approaches and future perspectives. Neuroimage 140:4–19. doi:10.1016/j.neuroimage.2016.02.012

Bergmann TO, Varatheeswaran R, Hanlon CA, Madsen KH, Thielscher A, Siebner HR. 2021. Concurrent TMS-fMRI for causal network perturbation and proof of target engagement. Neuroimage 237:118093. doi:10.1016/j.neuroimage.2021.118093

Bestmann S, Baudewig J, Frahm J. 2003. On the synchronization of transcranial magnetic stimulation and functional echo-planar imaging. Journal of Magnetic Resonance Imaging 17:309–316. doi:10.1002/jmri.10260

Betzel RF, Bassett DS. 2017. Multi-scale brain networks. Neuroimage 160:73–83. doi:10.1016/j.neuroimage.2016.11.006

Boonzaier J, Petrov PI, Otte WM, Smirnov N, Neggers SFW, Dijkhuizen RM. 2020. Design and evaluation of a rodent-specific transcranial magnetic stimulation coil: An in silico and in vivo validation study. Neuromodulation: Technology at the Neural Interface 23:324–334. doi:10.1111/ner.13025

Cash RFH, Weigand A, Zalesky A, Siddiqi SH, Downar J, Fitzgerald PB, Fox MD. 2021. Using brain imaging to improve spatial targeting of transcranial magnetic stimulation for depression. Biol Psychiatry 90:689–700. doi:10.1016/J.BIOPSYCH.2020.05.033

Crowther LJ, Porzig K, Hadimani RL, Brauer H, Jiles DC. 2013. Realistically modeled transcranial magnetic stimulation coils for lorentz force and stress calculations during MRI. IEEE Trans Magn 49:3426–3429. doi:10.1109/TMAG.2013.2247578

Dawson TM, Golde TE, Lagier-Tourenne C. 2018. Animal models of neurodegenerative diseases. Nat Neurosci 21:1370–1379. doi:10.1038/s41593-018-0236-8

El Arfani A, Parthoens J, Demuyser T, Servaes S, De Coninck M, De Deyn PP, Van Dam D, Wyckhuys T, Baeken C, Smolders I, Staelens S. 2017. Accelerated high-frequency repetitive transcranial magnetic stimulation enhances motor activity in rats. Neuroscience 347:103–110. doi:10.1016/j.neuroscience.2017.01.045

Fox PT, Narayana S, Tandon N, Sandoval H, Fox SP, Kochunov P, Lancaster JL. 2004. Column-based model of electric field excitation of cerebral cortex. Hum Brain Mapp 22:1–14. doi:10.1002/hbm.20006

Funke K. 2018. Transcranial magnetic stimulation of rodents Handbook of Behavioral Neuroscience. Elsevier. pp. 365–387. doi:10.1016/B978-0-12-812028-6.00020-3

Gordon EM, Chauvin RJ, Van AN, Rajesh A, Nielsen A, Newbold DJ, Lynch CJ, Seider NA, Krimmel SR, Scheidter KM, Monk J, Miller RL, Metoki A, Montez DF, Zheng A, Elbau I, Madison T, Nishino T, Myers MJ, Kaplan S, Badke D’Andrea C, Demeter D V., Feigelis M, Ramirez JSB, Xu T, Barch DM, Smyser CD, Rogers CE, Zimmermann J, Botteron KN, Pruett JR, Willie JT, Brunner P, Shimony JS, Kay BP, Marek S, Norris SA, Gratton C, Sylvester CM, Power JD, Liston C, Greene DJ, Roland JL, Petersen SE, Raichle ME, Laumann TO, Fair DA, Dosenbach NUF. 2023. A somato-cognitive action network alternates with effector regions in motor cortex. Nature 617:351–359. doi:10.1038/s41586-023-05964-2

Ilmoniemi RJ, Virtanen J, Ruohonen J, Karhu J, Aronen HJ, Näätänen R, Katila T. 1997. Neuronal responses to magnetic stimulation reveal cortical reactivity and connectivity. Neuroreport 8:3537– 3540. doi:10.1097/00001756-199711100-00024

Kätzel D, Zemelman B V, Buetfering C, Wölfel M, Miesenböck G. 2011. The columnar and laminar organization of inhibitory connections to neocortical excitatory cells. Nat Neurosci 14:100–107. doi:10.1038/nn.2687

Kinsler LE, Frey AR, Coppens AB, Sanders JV. 1999. Fundamentals of acoustics (4th Edition), John Wiley and Sons Inc.

Koponen LM, Nieminen JO, Ilmoniemi RJ. 2018. Multi-locus transcranial magnetic stimulation— theory and implementation. Brain Stimul 11:849–855. doi:10.1016/j.brs.2018.03.014

Koponen LM, Nieminen JO, Mutanen TP, Stenroos M, Ilmoniemi RJ. 2017. Coil optimisation for transcranial magnetic stimulation in realistic head geometry. Brain Stimul 10:795–805. doi:10.1016/j.brs.2017.04.001

Koponen LM, Stenroos M, Nieminen JO, Jokivarsi K, Gröhn O, Ilmoniemi RJ. 2020. Individual head models for estimating the TMS-induced electric field in rat brain. Sci Rep 10:17397. doi:10.1038/s41598-020-74431-z

Krieg TD, Salinas FS, Narayana S, Fox PT, Mogul DJ. 2013. PET-based confirmation of orientation sensitivity of TMS-induced cortical activation in humans. Brain Stimul 6:898–904. doi:10.1016/j.brs.2013.05.007

Lauer AM, El-Sharkawy A-MM, Kraitchman DL, Edelstein WA. 2012. MRI acoustic noise can harm experimental and companion animals. Journal of Magnetic Resonance Imaging 36:743–747. doi:10.1002/jmri.23653

Lefaucheur J-P, André-Obadia N, Antal A, Ayache SS, Baeken C, Benninger DH, Cantello RM, Cincotta M, de Carvalho M, De Ridder D, Devanne H, Di Lazzaro V, Filipović SR, Hummel FC, Jääskeläinen SK, Kimiskidis VK, Koch G, Langguth B, Nyffeler T, Oliviero A, Padberg F, Poulet E, Rossi S, Rossini PM, Rothwell JC, Schönfeldt-Lecuona C, Siebner HR, Slotema CW, Stagg CJ, Valls-Sole J, Ziemann U, Paulus W, Garcia-Larrea L. 2014. Evidence-based guidelines on the therapeutic use of repetitive transcranial magnetic stimulation (rTMS). Clinical Neurophysiology 125:1–57. doi:10.1016/j.clinph.2014.05.021

Lehto LJ, Canna A, Wu L, Sierra A, Zhurakovskaya E, Ma J, Pearce C, Shaio M, Filip P, Johnson MD, Low WC, Gröhn O, Tanila H, Mangia S, Michaeli S. 2020. Orientation selective deep brain stimulation of the subthalamic nucleus in rats. Neuroimage 213:116750. doi:10.1016/j.neuroimage.2020.116750

Li L, Yin Z, Huo X. 2007. The influence of low-frequency rTMS on EEG of rats. Neurosci Lett 412:143–147. doi:10.1016/j.neulet.2006.10.054

Lynn CW, Bassett DS. 2019. The physics of brain network structure, function and control. Nature Reviews Physics 1:318–332. doi:10.1038/s42254-019-0040-8

Massimini M, Ferrarelli F, Huber R, Esser SK, Singh H, Tononi G. 2005. Breakdown of cortical effective connectivity during sleep. Science (1979) **309**:2228–2232. doi:10.1126/science.1117256

Mengotti P, Käsbauer A-S, Fink GR, Vossel S. 2022. Combined TMS-fMRI reveals behavior-dependent network effects of right temporoparietal junction neurostimulation in an attentional belief updating task. Cerebral Cortex 32:4698–4714. doi:10.1093/cercor/bhab511

Mizutani-Tiebel Y, Tik M, Chang K-Y, Padberg F, Soldini A, Wilkinson Z, Voon CC, Bulubas L, Windischberger C, Keeser D. 2022. Concurrent TMS-fMRI: Technical challenges, developments, and overview of previous studies. Front Psychiatry 13:581. doi:10.3389/fpsyt.2022.825205

Navarro de Lara LI, Tik M, Woletz M, Frass-Kriegl R, Moser E, Laistler E, Windischberger C. 2017. High-sensitivity TMS/fMRI of the human motor cortex using a dedicated multichannel MR coil. Neuroimage 150:262–269. doi:10.1016/j.neuroimage.2017.02.062

Nestler EJ, Hyman SE. 2010. Animal models of neuropsychiatric disorders. Nat Neurosci 13:1161– 1169. doi:10.1038/nn.2647

Nieminen JO, Koponen LM, Ilmoniemi RJ. 2015. Experimental characterization of the electric field distribution induced by TMS devices. Brain Stimul 8:582–589. doi:10.1016/j.brs.2015.01.004

Nieminen JO, Koponen LM, Mäkelä N, Souza VH, Stenroos M, Ilmoniemi RJ. 2019. Short-interval intracortical inhibition in human primary motor cortex: A multi-locus transcranial magnetic stimulation study. Neuroimage 203:116194. doi:10.1016/j.neuroimage.2019.116194

Nieminen JO, Pospelov AS, Koponen LM, Yrjölä P, Shulga A, Khirug S, Rivera C. 2022a. Transcranial magnetic stimulation set-up for small animals. Front Neurosci 16. doi:10.3389/fnins.2022.935268

Nieminen JO, Sinisalo H, Souza VH, Malmi M, Yuryev M, Tervo AE, Stenroos M, Milardovich D, Korhonen JT, Koponen LM, Ilmoniemi RJ. 2022b. Multi-locus transcranial magnetic stimulation system for electronically targeted brain stimulation. Brain Stimul 15:116–124. doi:10.1016/j.brs.2021.11.014

Nurmi S, Karttunen J, Souza VH, Ilmoniemi RJ, Nieminen JO. 2021. Trade-off between stimulation focality and the number of coils in multi-locus transcranial magnetic stimulation. J Neural Eng 18:066003. doi:10.1088/1741-2552/ac3207

Nyrhinen MJ, Souza VH, Ilmoniemi RJ, Lin F-H. 2023. Acoustic noise generated by TMS in typical environment and inside an MRI scanner. SSRN. doi:10.2139/SSRN.4470011

Parthoens J, Wyckhuys T, Crevecoeur G, Stroobants S, Staelens S. 2012. Fast screening of transcranial magnetic stimulation paradigms in the rat using microPET. Journal of Nuclear Medicine 53.

Paus T, Jech R, Thompson CJ, Comeau R, Peters T, Evans AC. 1997. Transcranial magnetic stimulation during positron emission tomography: A new method for studying connectivity of the human cerebral cortex. The Journal of Neuroscience 17:3178–3184. doi:10.1523/JNEUROSCI.17-09-03178.1997

Rissanen IJ, Souza VH, Nieminen JO, Koponen LM, Ilmoniemi RJ. 2023. Advanced pipeline for designing multi-locus TMS coils with current density constraints. IEEE Trans Biomed Eng 70:2025– 2034. doi:10.1109/TBME.2023.3234119

Rosanova M, Casali A, Bellina V, Resta F, Mariotti M, Massimini M. 2009. Natural frequencies of human corticothalamic circuits. Journal of Neuroscience 29:7679–7685. doi:10.1523/JNEUROSCI.0445-09.2009

Rotenberg A, Muller P, Birnbaum D, Harrington M, Riviello JJ, Pascual-Leone A, Jensen FE. 2008. Seizure suppression by EEG-guided repetitive transcranial magnetic stimulation in the rat. Clinical Neurophysiology 119:2697–2702. doi:10.1016/j.clinph.2008.09.003

Rotenberg A, Muller PA, Vahabzadeh-Hagh AM, Navarro X, López-Vales R, Pascual-Leone A, Jensen F. 2010. Lateralization of forelimb motor evoked potentials by transcranial magnetic stimulation in rats. Clinical Neurophysiology 121:104–108. doi:10.1016/j.clinph.2009.09.008

Seewoo BJ, Etherington SJ, Feindel KW, Rodger J. 2018. Combined rTMS/fMRI Studies: An Overlooked Resource in Animal Models. Front Neurosci 12. doi:10.3389/fnins.2018.00180

Siddiqi SH, Kording KP, Parvizi J, Fox MD. 2022. Causal mapping of human brain function. Nat Rev Neurosci 23:361–375. doi:10.1038/s41583-022-00583-8

Siddiqi SH, Schaper FLWVJ, Horn A, Hsu J, Padmanabhan JL, Brodtmann A, Cash RFH, Corbetta M, Choi KS, Dougherty DD, Egorova N, Fitzgerald PB, George MS, Gozzi SA, Irmen F, Kuhn AA, Johnson KA, Naidech AM, Pascual-Leone A, Phan TG, Rouhl RPW, Taylor SF, Voss JL, Zalesky A, Grafman JH, Mayberg HS, Fox MD. 2021. Brain stimulation and brain lesions converge on common causal circuits in neuropsychiatric disease. Nat Hum Behav 5:1707–1716. doi:10.1038/s41562-021-01161-1

Siddiqi SH, Taylor JJ, Horn A, Fox MD. 2023. Bringing human brain connectomics to clinical practice in psychiatry. Biol Psychiatry 93:386–387. doi:10.1016/j.biopsych.2022.05.026

Siebner HR, Bergmann TO, Bestmann S, Massimini M, Johansen-Berg H, Mochizuki H, Bohning DE, Boorman ED, Groppa S, Miniussi C, Pascual-Leone A, Huber R, Taylor PCJ, Ilmoniemi RJ, de Gennaro L, Strafella AP, Kahkonen S, Kloppel S, Frisoni GB, George MS, Hallett M, Brandt SA, Rushworth MF, Ziemann U, Rothwell JC, Ward N, Cohen LG, Baudewig J, Paus T, Ugawa Y, Rossini PM. 2009. Consensus paper: Combining transcranial stimulation with neuroimaging. Brain Stimul 2:58–80. doi:10.1016/j.brs.2008.11.002

Slopsema JP, Peña E, Patriat R, Lehto LJ, Gröhn O, Mangia S, Harel N, Michaeli S, Johnson MD. 2018. Clinical deep brain stimulation strategies for orientation-selective pathway activation. J Neural Eng 15:056029. doi:10.1088/1741-2552/aad978

Souza VH, Nieminen JO, Tugin S, Koponen LM, Baffa O, Ilmoniemi RJ. 2022. TMS with fast and accurate electronic control: Measuring the orientation sensitivity of corticomotor pathways. Brain Stimul 15:306–315. doi:10.1016/J.BRS.2022.01.009

Surendrakumar S, Rabelo TK, Campos ACP, Mollica A, Abrahao A, Lipsman N, Burke MJ, Hamani C. 2023. Neuromodulation therapies in pre-clinical models of traumatic brain injury: Systematic review and translational applications. J Neurotrauma 40:435–448. doi:10.1089/neu.2022.0286

Tang A, Thickbroom G, Rodger J. 2017. Repetitive transcranial magnetic stimulation of the brain. The Neuroscientist 23:82–94. doi:10.1177/1073858415618897

Tang W, Choi EY, Heilbronner SR, Haber SN. 2020. Nonhuman primate meso-circuitry data: a translational tool to understand brain networks across species. Brain Structure and Function 2020 226:1 226:1–11. doi:10.1007/S00429-020-02133-3

Tardelli GP, Souza VH, Matsuda RH, Garcia MAC, Novikov PA, Nazarova MA, Baffa O. 2022. Forearm and hand muscles exhibit high coactivation and overlapping of cortical motor representations. Brain Topogr 35:322–336. doi:10.1007/s10548-022-00893-1

Tastevin M, Richieri R, Boyer L, Fond G, Lançon C, Guedj E. 2020. Brain PET metabolic substrate of TMS response in pharmaco-resistant depression. Brain Stimul 13:683–685. doi:10.1016/j.brs.2020.02.014

Tik M, Woletz M, Schuler A-L, Vasileiadi M, Cash RFH, Zalesky A, Lamm C, Windischberger C. 2023. Acute TMS/fMRI response explains offline TMS network effects – An interleaved TMS-fMRI study. Neuroimage 267:119833. doi:10.1016/j.neuroimage.2022.119833

Tugin S, Souza VH, Nazarova MA, Novikov PA, Tervo AE, Nieminen JO, Lioumis P, Ziemann U, Nikulin V V., Ilmoniemi RJ. 2021. Effect of stimulus orientation and intensity on short-interval intracortical inhibition (SICI) and facilitation (SICF): A multi-channel transcranial magnetic stimulation study. PLoS One 16:e0257554. doi:10.1371/journal.pone.0257554

Turner JG, Bauer CA, Rybak LP. 2007. Noise in animal facilities: why it matters. J Am Assoc Lab Anim Sci 46:10–3.

Weise K, Numssen O, Kalloch B, Zier AL, Thielscher A, Haueisen J, Hartwigsen G, Knösche TR. 2023a. Precise motor mapping with transcranial magnetic stimulation. Nat Protoc 18:293–318. doi:10.1038/s41596-022-00776-6

Weise K, Worbs T, Kalloch B, Souza VH, Jaquier AT, Van Geit W, Thielscher A, Knösche TR. 2023b. Directional sensitivity of cortical neurons towards TMS-induced electric fields. Imaging Neuroscience 1:1–22. doi:10.1162/imag_a_00036

Wu T, Fan J, Lee KS, Li X. 2016. Cortical neuron activation induced by electromagnetic stimulation: a quantitative analysis via modelling and simulation. J Comput Neurosci 40:51–64. doi:10.1007/s10827-015-0585-1

Wyckhuys T, De Geeter N, Crevecoeur G, Stroobants S, Staelens S. 2013. Quantifying the effect of repetitive transcranial magnetic stimulation in the rat brain by μsPECT CBF scans. Brain Stimul 6:554–562. doi:10.1016/J.BRS.2012.10.004

Yang Y, Qiao S, Sani OG, Sedillo JI, Ferrentino B, Pesaran B, Shanechi MM. 2021. Modelling and prediction of the dynamic responses of large-scale brain networks during direct electrical stimulation. Nat Biomed Eng 5:324–345. doi:10.1038/s41551-020-00666-w

