## Supplementary Material for "A multi-channel TMS system enabling accurate stimulus orientation control during concurrent ultra-high-field MRI for preclinical applications"

### Multi-channel TMS power electronics and connectivity

The multi-channel transcranial magnetic stimulation (mTMS) system was designed for safe use inside the shielded room of a 9.4-T magnetic resonance imaging (MRI) scanner for small animals. Below, we described safety features in addition to those presented in the main text:

- The power electronics cabinet includes an electronic safety system that immediately discharges the pulse capacitors if the cabinet door is opened and has an external emergency stop button.
- With the power electronics inside the shielded room, we reduce the length of the cables connecting to the transducer minimizing undesirable inductance and resistance critical for the system operation.

- The metal structure of the cabinet is earthed with a separate cable, which remains connected even when the power cable is disconnected. The earthing follows a "star-ground" design, thus eliminating potential ground loops.
- The electronics are isolated by an isolation transformer (resistance to ground >2 MΩ), providing added user safety. There are no electrical connections between the power electronics and operator control cabinets.
- For thermal management, three 120 mm diameter fans push out air from the top of the cabinet, and the replacement air is mainly taken in from around the cabinet's front door, providing airflow from front to back and bottom to top.
- A custom-made door handle, lock, and non-magnetic copper keys were manufactured to prevent unauthorized access to the cabinet. The system also contains an emergency stop button.
- The only major magnetizable component in the system that could not be placed outside the shielded room was a 2000 VA isolation transformer.
- Whenever possible, internal components were fixed to the cabinet's structure.
- In the connection box, the coil cables are connected with lugs to a 2.5-m-long low-inductance TMS coil cable (The Magstim Company, Ltd., UK) that is connected in series with another 2.5-m-long low-inductance TMS cable (Nexstim, Plc., Finland) connecting to the mTMS power cabinet through a medical-grade connector (Nexstim).
- The mTMS system supports connection to digital temperature sensors that can be embedded into the transducer to monitor its temperature, and the signal path protects against coil-induced overvoltage.
- As only optical signals are transferred through the waveguide, external electrical and RF noise coupling to the MRI scanner has been eliminated.

**mTMS transducer design, manufacturing, and calibration**

Additional features and methods descriptions for the design, manufacturing, and calibration of the mTMS transducer are listed below.

The inner base of the outer case contains small studs, and the bottom coil plate has matching cavities. While the transducer is resting on the studs, a 1-mm air gap is present between the enclosure and the coil plates, reducing the mechanical coupling between them and providing an additional safety barrier in case of fractures in the coil plates.

The duration of the trapezoidal monophasic pulse waveform delivered by the mTMS power electronics was customized for each TMS coil based on measurements with a Rogowski probe (CWT 60B, Power Electronic Measurements, Ltd., UK) connected to an oscilloscope (InfiniiVision MSOX3034T, Keysight Technologies, Inc., USA) to ensure that no current was left circulating in the mTMS system after a pulse.

The self-inductance of the mTMS coils was measured with an LCR meter (1-kHz reference frequency; ELC-130, Escort Instruments Corp., Taiwan) and the resistance with a 4-wire measurement setup using a benchtop multimeter (HP 34401A; Hewlett-Packard Company, USA).

**mTMS pulse acoustic noise**

The beginning of the far field is given roughly by $L^2 f/4\mathrm{v}$, where $L$ is the largest dimension of the source, $f$ is the frequency, and v is the speed of sound [1]. For a TMS pulse, the far field starts at about 60 cm away from the coil center.

The acoustic noise reduction was calculated by measuring the SPL for all E-field orientations at 20% MSO stimulation intensity with and without the earplug. The average reduction between these two conditions was added to the SPL measurements utilizing the earplug.

Considering the rat's safety, we reported all SPL values without any weighting (Z-weighting); thus, our results may slightly overestimate the acoustic noise humans perceive by about 2 dB compared to human audiograms (C- and A-weightings). Previous studies have addressed the sensitivity of laboratory animals, more specifically rats, to sound exposure [2–5].

Impulse sounds of 160 dB in the human-perceivable auditory range (peaking at around 4 kHz) might also cause temporal and permanent hearing threshold changes in different species, such as guinea pigs, rats, and mice [2].

**Supplementary References**


[1] Kinsler LE, Frey AR, Coppens AB, Sanders JV. Fundamentals of acoustics (4th Edition). 1999.

[2] Duan M, Laurell G, Qiu J, Borg E. Susceptibility to impulse noise trauma in different species: guinea pig, rat and mouse. Acta Otolaryngol 2008;128:277–83. https://doi.org/10.1080/00016480701509941.

[3] Holt AG, Kühl A, Braun RD, Altschuler R. The rat as a model for studying noise injury and otoprotection. J Acoust Soc Am 2019;146:3681–91. https://doi.org/10.1121/1.5131344.

[4] Turner JG, Bauer CA, Rybak LP. Noise in animal facilities: why it matters. J Am Assoc Lab Anim Sci 2007;46:10–3.

[5] Lauer AM, El-Sharkawy A-MM, Kraitchman DL, Edelstein WA. MRI acoustic noise can harm experimental and companion animals. Journal of Magnetic Resonance Imaging 2012;36:743–7. https://doi.org/10.1002/jmri.23653.
